# Protein Barcoding and Next-Generation Protein Sequencing for Multiplexed Protein Selection, Analysis, and Tracking

**DOI:** 10.1101/2024.12.31.630920

**Authors:** Mathivanan Chinnaraj, Haidong Huang, Sebastian Hutchinson, Michael Meyer, Douglas Pike, Marco Ribezzi, Sharmin Sultana, Ghada H. Mansour, Derrek Ocampo, Fengling Ding, Meredith L. Carpenter, Ilya Chorny, John Vieceli

## Abstract

Protein barcoding has emerged as a powerful tool for the multiplexed identification and characterization of proteins, providing a mechanism for precise tracking of protein affinity, location, and expression. In this study, we describe the development of a protein barcoding workflow for use with single-molecule Next-Generation Protein Sequencing™ (NGPS™) on the benchtop Platinum^®^ and Platinum^®^ Pro instruments. We present data on the validation of eight peptide barcodes, each designed to minimize detection bias and maximize sensitivity across various experimental conditions. We have also optimized the design of expression constructs to decrease both the hands-on time and input requirements of the workflow. In this workflow, affinity-tagged proteins are expressed with unique peptide barcodes. Following experimental selection or treatments, the proteins are purified, and the peptide barcodes are cleaved and sequenced on the Platinum instrument. We demonstrate that we can detect barcodes at 400 fmol of sample input concentration within the eight-plex mixture, and at 50 fmol of sample input for individual barcodes. We also show the capacity of this barcoding approach to achieve a ten-fold dynamic range, underscoring its sensitivity in recovering variants with low abundance. Through the combination of protein barcoding and NGPS, we lay the groundwork for future studies aimed at characterizing protein interactions and improving targeted drug delivery strategies.

## Introduction

In recent years, protein/peptide barcoding has gained attention as a powerful method for advancing protein analysis^1-9^. This approach leverages the unique ability of short peptide sequences to encode information, providing an efficient and flexible means of tracking and characterizing proteins. Unlike traditional labeling techniques, peptide barcodes can be easily genetically encoded, offering a straightforward way to label proteins within complex biological systems without disrupting their native function. This versatility has made protein barcoding an increasingly valuable tool in proteomics and functional genomics, enabling more precise studies of protein behavior and interactions in a variety of experimental contexts^1,3-10^.

Protein barcodes have already been developed and applied in a variety of settings, leveraging the use of mass spectrometry for detection and decoding. For instance, “flycodes” have been used in nanobody screening to rapidly assess protein interactions^3^, and abiotic peptides have been employed for large-scale screening of small molecule libraries^2^. Despite these advances, several challenges remain, particularly in the ability to directly read protein barcode sequences with quantitative accuracy and single-molecule resolution. Ionization efficiency can vary between different peptide sequences, and signal overlap can complicate interpretation^11^. Furthermore, mass spectrometry requires expensive equipment and extensive expertise to generate and analyze data. This gap has hindered the broader application of peptide barcoding in proteomics and functional screening.

Recent innovations in single-molecule protein sequencing may offer a solution to these limitations. Novel protein sequencing technologies, including the Platinum^®^ and Platinum Pro^®^ instruments, allow for the direct sequencing of protein barcodes with single-molecule resolution and an accessible benchtop workflow^12^. NGPS on Platinum involves the use of fluorescently tagged N-terminal amino acid (NAA) recognizer proteins to determine the order of amino acids in a peptide bound to a semiconductor chip^12^ (**Figure 1A**). By distinguishing peptides based on their amino acid sequences rather than mass/charge ratios, NGPS overcomes some of the key challenges of mass spectrometry, such as the inability to resolve peptides with identical or highly similar amino acid compositions^13^. This capability enables precise identification of protein sequences and opens the door to a range of new applications in protein characterization. In addition, the straightforward sample preparation and data analysis workflows make NGPS a highly accessible approach to protein barcode implementation.

**Figure 1.**
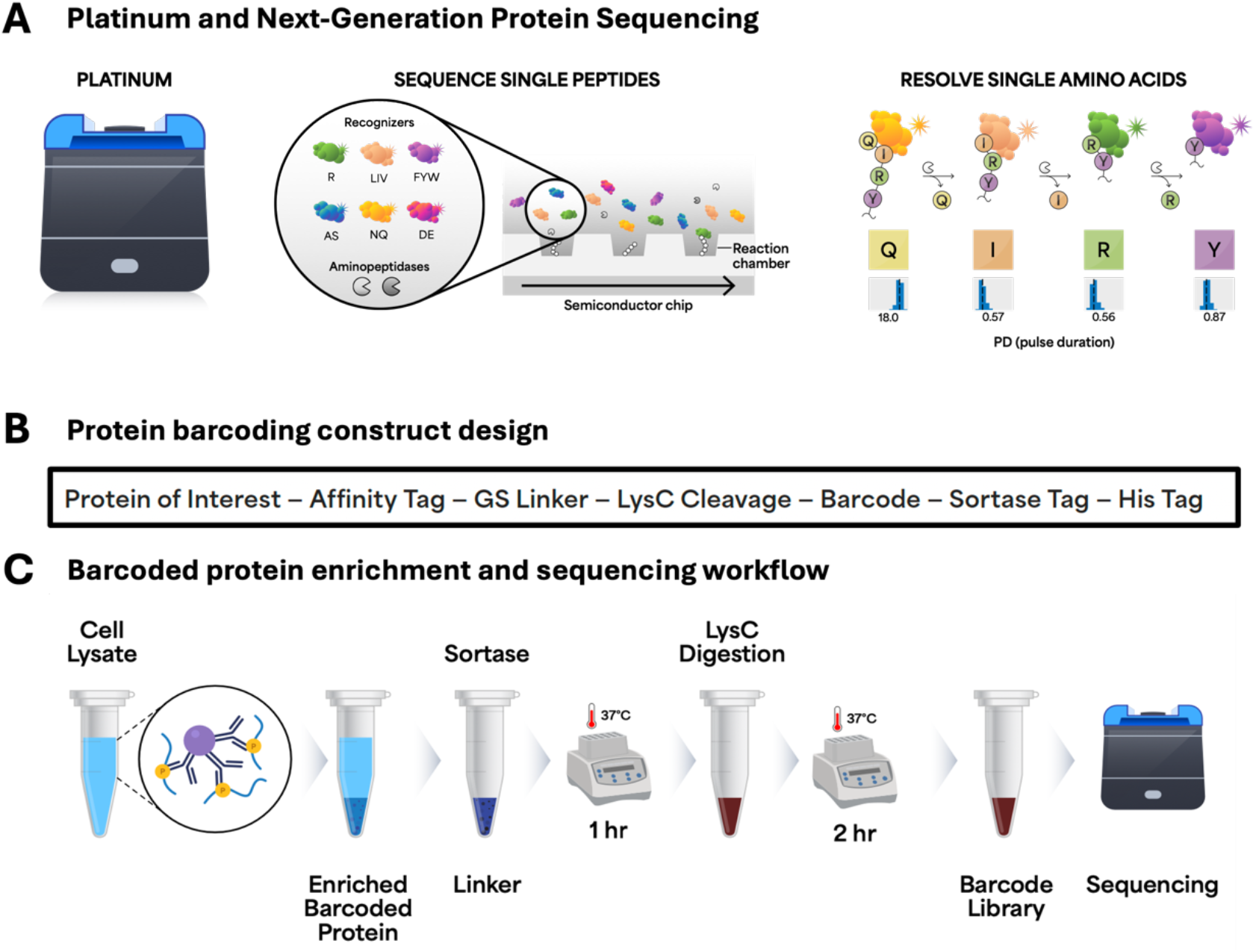
Overview of Platinum sequencing, protein barcoding construct design, and barcoding workflow. A) Overview of the Platinum instrument and the principle of Next-Generation Protein Sequencing. After single peptides are bound to the semiconductor chip, fluorescently tagged amino acid recognizers (six recognizers for 13 amino acids) bind each N-terminal amino acid. After aminopeptidase cleavage, the next amino acid is bound. B) Barcoding construct design includes the protein of interest, followed by an affinity tag for purification, a short linker, a LysC cleavage site, the peptide barcode, a sortase tag for attachment of a covalent linker for sequencing on Platinum, and an optional His tag for purification. C) Barcoded protein enrichment and barcode sequencing workflow showing the steps going from cell lysate to sequencing.

The concept of protein barcoding is rooted in the success of DNA barcoding, a technique that has been widely applied in genomics and transcriptomics. DNA barcodes are short sequences of DNA that encode information and can be efficiently decoded using next-generation sequencing. This approach enables high-throughput analyses such as tracking sample identity in multiplexed libraries and mapping single-cell gene expression^14-19^. However, while DNA barcodes have found broad use in molecular biology, their application to protein analysis has been more limited due to the need to retain a genotype-phenotype connection for readout, as well as the inability to directly detect successful translation with DNA barcodes^1,3,8,9^.

One area where protein barcoding has shown particular promise is in the development of nucleic acid therapies ^4,5,7^. For instance, nucleic acid delivery systems, such as lipid nanoparticles (LNPs), often require tracking of both the uptake and functional delivery of therapeutic cargo to specific tissues or cells. While DNA barcodes have been used to track LNP uptake, they can fail to confirm the functional delivery and activity of the encoded proteins ^4,5,7,14^. Protein barcodes, on the other hand, can provide direct readouts of protein function and localization, offering a more precise and scalable method for tracking the success of nucleic acid delivery vectors^5,7^. In protein engineering, protein barcodes also hold significant potential. By tagging different variants of peptides with unique sequences, researchers can use barcoding to track the functional properties of engineered proteins in complex screening assays^1,3,6,8^. This approach enables the rapid identification of proteins with desirable traits, such as improved stability, binding affinity, or enzymatic activity, which are critical for the development of new biotherapeutics.

In addition to gene therapy and protein engineering, protein barcoding has applications in other areas, such as studying protein-protein interactions, tracking protein subcellular localization, and even screening small-molecule libraries ^1-3,6,8^. The ability to encode functional information within peptides and decode it with high accuracy and resolution will enable researchers to gain deeper insights into complex cellular biology.

In this study, we developed a protein barcoding workflow combined with NGPS as a tool for advancing protein characterization with an accessible benchtop workflow. We then evaluated key performance metrics, including dynamic range and limit of detection, in the context of an optimized set of eight barcodes. This study serves as a foundation for the implementation of protein barcoding (now commercially available in the Barcoding Kit from Quantum-Si) and NGPS workflows across a range of applications.

## Experimental Procedures

### Barcode design and optimization

To design barcodes compatible with the Platinum sequencing and analysis platform, we iteratively refined an initial large set of candidate sequences. First, we generated *recognizer-ordered sequences (ROS)* by assigning each amino acid recognizer a unique symbol (e.g., “1” for the Arginine Recognizer) and ensuring that all six recognizers in the V3 Sequencing Kit (**Figure 1A**) were included, with no two consecutive recognizers being the same. We then expanded these ROSs into full amino acid barcode sequences by enumerating all valid residue substitutions for each recognizer, evaluating each candidate’s predicted performance using a kinetic database of pulse durations, and discarding any prone to dropout (**Figure 2A**).

**Figure 2.**
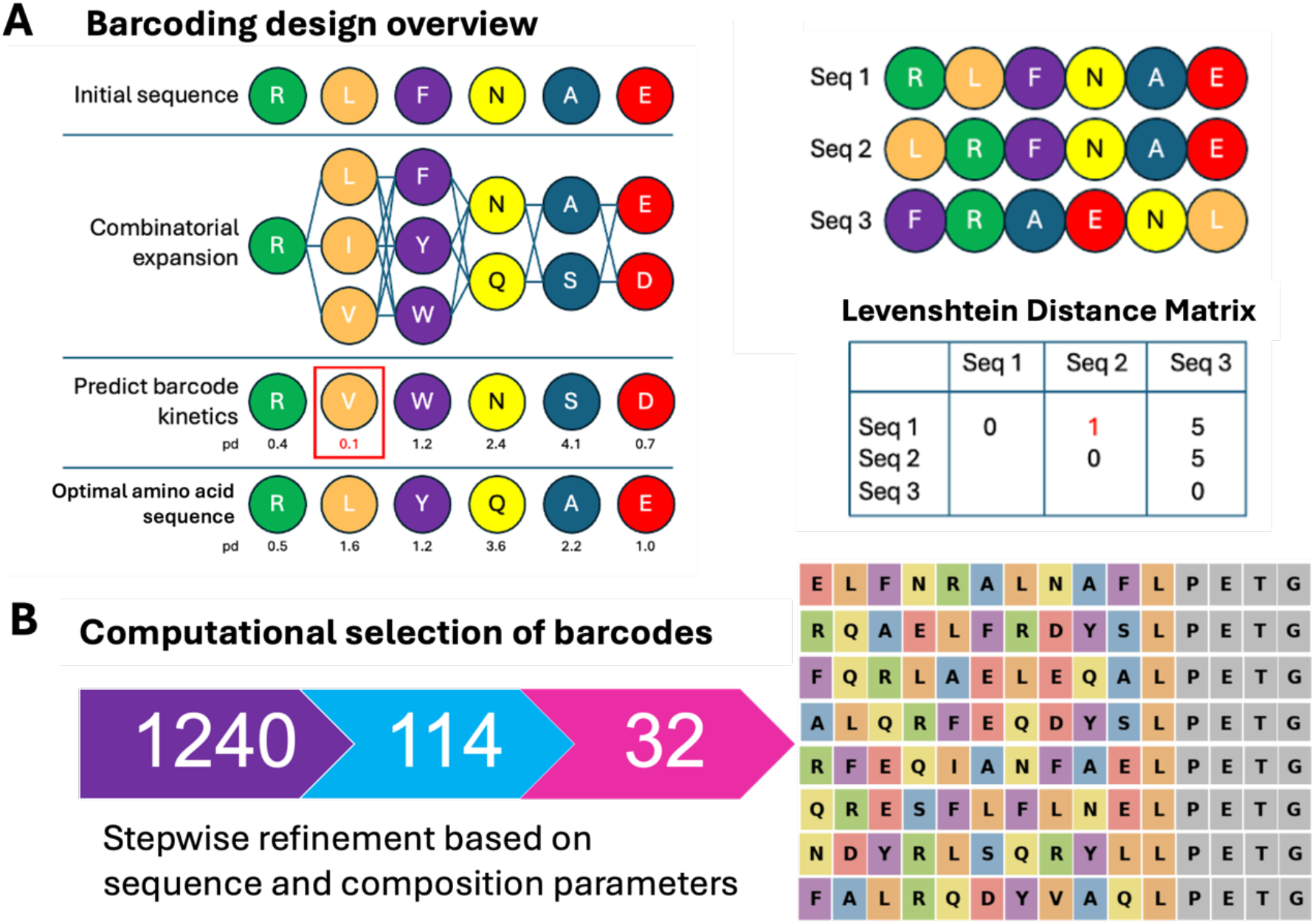
Computational design of protein barcodes for NGPS. A) Barcode design workflow selects optimal barcode designs by taking into account protein sequencing kinetics and Levenshtein edit distance to produce barcodes with optimal properties for multiplexing. B) Schematic of the computational selection and refinement of barcodes to the eight used in this study.

To ensure reliability despite potential errors (e.g., missed or substituted residues), we calculated the Levenshtein distance between ROS and required a minimum distance between every pair. This ensured each barcode remained uniquely identifiable, even if partial errors occurred (**Figure 2A**).

To compute error-resistant barcode sets, we employed a heuristic approach. We created an empty barcode set, then randomized all candidate ROS and iterated over each ROS in this pool, extending the barcode set only if the new candidate met the edit-distance threshold. This process was repeated 1,000 times.

From the resulting population of candidate sets, we chose the one best satisfying both size and composition criteria (**Figure 2B**). This yielded barcode sets with strong error tolerance and high confidence in their unique identification.

### Construct Design and Protein Purification

The following barcodes were designed and both 1) added to the full-length protein construct as well as 2) produced as synthetic barcodes:

BC028, DYKDDDDKGGGGSGGGGSKRFEQIANFAELPETGH;

BC032, DYKDDDDKGGGGSGGGGSKRQAELFRDYSLPETGH;

BC049, DYKDDDDKGGGGSGGGGSKFQRLAELEQALPETGH;

BC051, DYKDDDDKGGGGSGGGGSKFALRQDYVAQLPETGH;

BC067, DYKDDDDKGGGGSGGGGSKQRESFLFLNELPETGH;

BC075, DYKDDDDKGGGGSGGGGSKNDYRLSQRYLLPETGH;

BC079, DYKDDDDKGGGGSGGGGSKALQRFEQDYSLPETGH;

BC096, DYKDDDDKGGGGSGGGGSKELFNRALNAFLPETGH

The synthetic barcodes were custom synthesized by InnoPep (San Diego, CA), each supplied at 3 mg and with a purity greater than 95%. All synthetic peptides featured an N-terminal H and C-terminal carboxylic acid block NH2. They were initially reconstituted in DMSO to a concentration of 10 mM and stored at –20°C until ready for the barcoding kit workflow. The peptides then go through the same sample preparation steps as the purified protein (below and **Figure 1C)**.

The five full-length proteins (IFNg-BC032, PTEN-BC049, TAU441-BC051, UCHL1-BC075, and p53-BC096) were cloned in pET21(a) with (i) c-terminal FLAG tag for affinity purification, (ii) a flexible GS linker as a spacer between affinity tag and barcode, (iii) LysC-cleavage site, (iv) peptide barcode, (v) sortase tag, and an optional (vi) 6x His-tag. See **Figure 1B** for an overview of the final construct design. All vectors were transformed into *E. coli* strain BL21(DE3) (Genscript, New Jersey, USA) to express in Super Broth Auto-Induction Media (Grisp Research Solutions, Portugal) at 37°C, then transferred into 18°C for overnight shaking at 200 RPM. The purification was done with anti-FLAG antibody magnetic beads to selectively capture FLAG-tagged, barcoded proteins of interest using either Pierce™ Anti-DYKDDDDK Magnetic Agarose (ThermoFisher; Cat. No. A36797) or Anti-FLAG® M2 Magnetic Beads (MilliporeSigma; Cat. No. M8823). The optional primary or secondary purification was done using cobalt-based IMAC Talon Superflow (Cytiva, USA) resin. Enriched protein was buffer exchanged in 50 mM Tris-HCL pH 7.5, and 150 mM NaCl to be compatible with sortase reactions, and the concentration of each protein was quantified using A280 Nanodrop Spectrophotometer (Thermo Fisher Scientific). The full sequences of all five proteins are shown in **Table S1A**.

Additionally, for the initial study (See Workflow section below) we also designed a synthetic peptide (BC265, DYKDDDDKGGGGSGGGGSKALQFRLFHTDDDLPETGH) and a version that lacked the GS linker and the lysine cleavage site between the GS linker and barcode (BC228, DYKDDDDKALQFRLFHTDDDLPETGH). We also designed two protein constructs: SARS-CoV2-S1-RBD domain (R319-F541) protein with FLAG tag, barcode sequence (ALQFRLFHTDDD), sortase tag, and optional 6xHis-tag was cloned into pcDNA 3.1 vector and expressed in HEK293 as a secreted protein. The full-length p53 protein with FLAG tag, barcode sequence (LFQARLFHTDDD), sortase tag, and optional 6xHis-tag was cloned into pET21 and expressed in *E. coli* by BPS Biosciences (San Diego, CA). The full sequences of these two proteins are shown in **Table S1B**.

### G-linker Production

A peptide-DNA-streptavidin conjugate was used as the linker to position barcode peptides on the chip surface. A DNA duplex was used as the structural scaffold to keep peptides away from the surface matrix. A fluorescent dye was conjugated to one end of the DNA with an amino modifier near the Streptavidin for loading quantification. The other end of DNA was modified with an O2’-propargyl adenosine as the conjugation handle for an aspartate-rich peptide spacer. The N-terminus of the aspartate-rich peptide is modified with a polyG moiety as the sortase conjugation handle. The identity of the polyG-peptide-DNA-streptavidin conjugate (G-linker) was confirmed by SEC-MS on an Agilent QTOF system.

### Workflow (Enrichment, Ligation, Cleavage) Development

We carried out two different workflows through the course of the study. In the first version, Workflow A (**Figure S1A**), the protein is enriched via affinity tag using anti-FLAG antibody magnetic beads at a minimum sample input of 500 pmol. We then performed the sortase ligation reaction with Picolyl-Azide-Gly-Gly-Gly (Vector labs, USA) at 37°C for 1 hr; this reaction results in covalent attachment of barcoded protein or peptides to an azide handle. After washing away excess Gly-Gly-Gly-Picolyl-Azide, we then added K-Linker (Quantum-Si, USA). The barcode-ligated azide handle and DBCO moiety on the K-Linker were covalently attached via Strain-Promoted Alkyne-Azide Cycloaddition (SPAAC) click reaction at 37°C for 16 hours, then the excess K-Linker was washed away. Finally, barcode linked K-Linker was cleaved from protein using enterokinase (Invitrogen, USA) or LysC enzymes (Quantum-Si, USA) at 37°C for 2 hours or longer. The prepared barcode libraries were then loaded and sequenced on the Platinum instrument.

In the second version, Workflow B (**Figure 1C, and Figure S1B**), the protein is enriched via affinity tag using anti-FLAG antibody magnetic beads at a sample input of 50 fmol or higher. We then incubate with 100 nM G-linker and 2 uM Sortase A5 enzyme (Quantum-Si, USA) in sortase reaction buffer (50 mM Tris-HCL pH 7.5, 150 mM NaCl, and 5 mM CaCl_2_) at 37°C for 1 hr on thermomixer at 1000 RPM. This reaction results in covalent attachment of the G-linker to the barcoded protein, eliminating the need for click reactions from workflow A and reducing the required sample input 10,000-fold. Finally, the G-linker ligated barcode was cleaved from protein using LysC enzyme at 37°C for 2 hours on thermomixer at 1000 RPM. This step releases the barcode-ligated G-linker from the FLAG-enriched protein of interest or peptides still bound on beads. The G-linker allows direct and stable anchoring of barcodes to the semiconductor chip surface. The ligated barcode libraries were stored at –20°C until sequencing.

### Barcode Sequencing on Platinum

The sequencing of the barcodes was carried out on a Quantum-Si Platinum instrument according to the manufacturer’s instructions. Briefly, approximately 100 pM of the barcoded G-linker was loaded, followed by the removal of excess, unbound barcodes. All sequencing was performed with the Sequencing Kit V3 (https://www.quantum-si.com/resources/product-data-sheets/platinum-instrument-and-sequencing-kit-v3-data-sheet/), which includes N-terminal amino acid (NAA) recognizers for 13 of the 20 canonical amino acids. Specifically, the kit contains a set of six NAA recognizers for LIV, FYW, and R, as previously described^12^, along with additional recognizers for AS, DE, and NQ (**Figure 1A)**. The binding and dissociation of these NAA recognizers to the immobilized peptide barcodes are monitored in real time as individual on-off events. NAAs from immobilized peptides are sequentially cleaved by aminopeptidases, allowing the next amino acid to be exposed for NAA recognizers to bind (**Figure 1A)**. This process is repeated throughout the 10-hour run time.

### Data Analysis

The Platinum instrument produces pulse calls as output of the raw sequencing data during real-time data collection. The pulse calls were transferred to the Platinum Analysis Software. Initially, all runs were analyzed using the Primary Analysis v2.8.0, which produces recognition segments of detected regions of interest at the aperture level. Then all runs go through secondary analysis using the Peptide Alignment v2.9.0, which takes primary analysis as an input and aligns observed recognition segments to the barcode reference at the aperture level. The resulting aperture-level results are filtered with a threshold score of 4.0 or above, then False Discovery Rate (FDR) is calculated with 20 decoy peptides and a reverse sequence of the reference. In general, an FDR of 10% or lower is required for a positive identification of barcodes. The number of apertures that pass strict filtering, FDR, and alignment are all grouped per barcode to plot the total number of alignments per run and total number of alignments for each barcode. Mean FDR is also calculated per identified barcode.

### Mean Absolute Percent Error

Mean Absolute Percent Error (MAPE) was computed for each experiment. For each barcode, percent error was computed by taking the absolute value of the predicted fraction minus the known fraction in the sample, and that result was divided by the true fraction. The mean of individual barcode percent errors across all samples is reported as the MAPE.

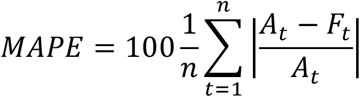

## Results

### Barcode construct design and testing

As a first step in this study, we set out to design and test expression constructs for barcoded proteins. To achieve efficient enrichment of barcoded protein expressed in cell or tissue, we designed constructs containing a FLAG tag and a unique barcode sequence, followed by a sortase tag with an optional 6xHisTag (**Figure 1B)**. We selected the FLAG affinity tag for several reasons: 1) it enables enrichment down to 15 fmol input from cell or tissue lysate; 2) it is easily accessible on the surface of the protein due to its charged residues and hydrophilic nature; 3) its smaller footprint reduces folding issues usually associated with larger affinity tags on smaller proteins; and 4) it can easily be cleaved by endopeptidase enterokinase (enteropeptidases), which recognizes DDDDK of the FLAG affinity handle and digests C-terminally to K. We also added a sortase tag as part of every barcode construct design to allow specific covalent modification to the barcode attached to the protein. Sortase A Pentamutant, an enzyme, is an engineered version of the wild-type sortase from *Staphylococcus aureus* that shows significantly higher activity than the wild-type sortase ^20^. Sortase belongs to a class of transpeptidases that utilize an active site cysteine thiol to modify proteins by recognizing and cleaving a carboxy-terminal sorting signal, LPXTG (where X is any amino acid), between the threonine and glycine residues. A nucleophile-containing poly-glycine sequence, (Gly)n (where n = 3 or more glycine residues), is used to attach a wide variety of labels such as peptides, DNA, carbohydrates, or fluorophores.

For the initial testing of this approach, we generated and loaded the following barcoded proteins on FLAG antibody beads: a synthetic peptide BC228, SARS-CoV2-S1-RBD, and p53. We then followed Workflow A as described in the **Experimental Procedures** section and shown in **Figure S1A**. The prepared libraries were then sequenced on Platinum. These steps resulted in successful sequencing, as shown in **Figure S2A-C**; however, the sample input was 500 pmol and the overall reaction time was 2 days. Thus, we focused on reducing the time and input requirements. We first created a unique G-linker, which contains polyG as a nucleophile for a sortase-mediated ligation (**Figure 1B-C** and **Figure S1B)**. Elimination of the DBCO click reactions from the K-Linker allowed us to attach the barcode directly and load it onto the chip for sequencing. However, this introduces another issue, as the enterokinase has promiscuity with the G-linker, and it also has difficulty accessing the cleavage site while the FLAG antibody beads are bound to the FLAG tag on the barcoded protein. To eliminate these issues, a flexible GS Linker (GGGGSGGGGS) was added between the affinity handle and barcode sequence (**Figure 1B-C** and **Figure S1B)**. An additional amino acid, lysine (K), was also added between the spacer and N-terminus of the barcode sequence (e.g. BC265) to replace the enterokinase with LysC as a cleavage protease. LysC has no promiscuity with the G-linker, and LysC enzymatic cleavage separates the barcode from the FLAG-captured protein. The flexible GS Linker helps create a spacer for easy accessibility of affinity enrichment, allows flexible folding, and its hydrophilic nature helps keep the LysC cleavage site on the protein surface for easy accessibility.

The combination of these unique tags, including the barcode, comprises less than 35 amino acids in length, minimizing structural folding complications arising from larger and bulky tags. This modified workflow also enables faster enrichment of barcodes from cell lysate to sequencing. Overall, these design changes with the newly created G-linker workflow as shown in **Figure 1B-C** and **Figure S1B** resulted in unprecedented sensitivity, enabling a 10,000-fold reduction in sample input from 500 pmol down to 50 fmol (**Figure S1B, Figure S2D**). Furthermore, the total time from cell lysate to loading on chip was reduced from two days to less than six hours, with less than one hour of hands-on time.

Following successful optimization of the workflow, we next refined the process for computational generation of barcodes (**Figure 2**). The barcodes are a unique sequence of 10 to 12 amino acids that are optimized for NGPS. We generated over a thousand barcodes, with each set containing 114 barcodes with equal sequencing capabilities, reduced bias, and low confusability between sequences, allowing random combination of any barcodes within a given pool (**Figure 2B)**. For initial validation, we selected a set of eight peptide barcode sequences optimized for Platinum sequencing that reliably produce distinct sets of barcodes with minimal false discovery rates (FDR) (see **Experimental Procedures** and **Figure S3A-B**). These eight barcodes are shown in **Table 1**, and the resulting sequencing kinetics summary for each barcode is shown in **Figure S4A-H**.

**Table 1.**
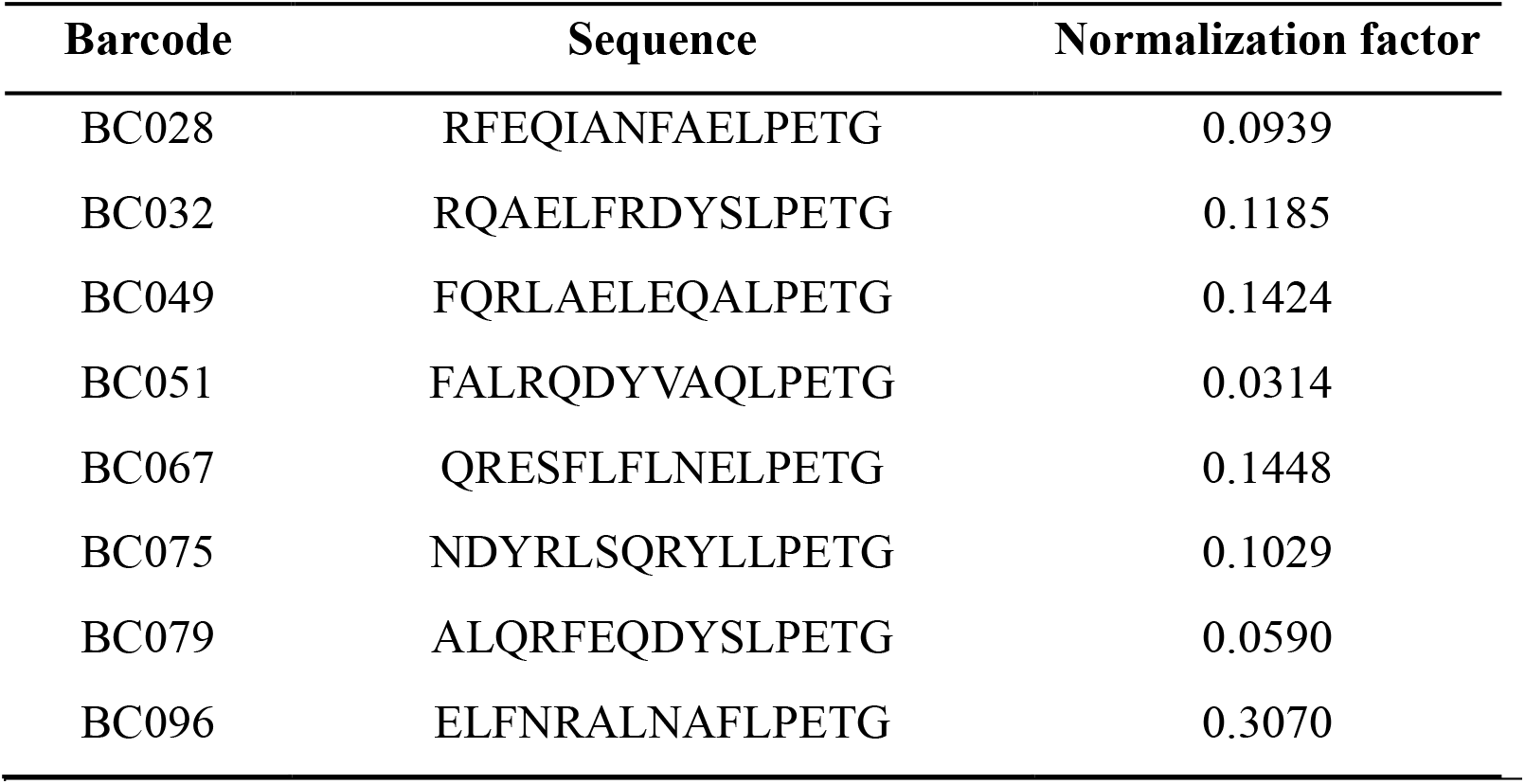
Summary of normalization factors used for each barcode.

### Normalization of barcodes in mixtures

After selecting these barcodes, we then sought to derive a set of normalization factors to increase linearity and reduce bias in multiplex mixtures. We mixed all eight barcodes at equimolar concentration to produce 1:1 mixture of plexity of eight each at 3.125 pmol (62.5 nM), with total sample input of 25 pmol. The normalization factors were initially generated by performing over 25 sequencing runs, resulting in over 200 data points from 1:1 mix, 10-fold, and 100-fold dynamic range mixtures of eight barcodes (**Figure S5)**. Runs were repeated in triplicate and with loading at 33 pM, 100 pM, and 300 pM. To calculate the normalization factors, we took the raw alignments for each barcode on each run and divided by total alignments to generate raw observed fractions. These raw observed fractions were re-normalized by known expected fractions, resulting in a pre-normalization factor. The median pre-normalization factor was taken to re-normalize, generating final normalization factors as shown **Table 1** and **Figure S5**.

### Normalization and reproducibility in 8-barcode mixtures

We performed an additional eight runs of 1:1 equimolar mix of all eight barcodes at 25 pmol total sample input and then applied the above established normalization factors to extract relative abundance of each barcode. As shown in **Figure 3A**, the alignments were converted to normalized alignments by dividing the normalization factor for each barcode, then each of the normalized alignments was divided by the sum of normalized alignments to extract the relative fraction of each observed barcode. The cumulative plot of normalized alignments for each barcode across eight runs is shown in **Figure 3B**. All eight barcodes were successfully identified with an FDR below the 10% cutoff **(Figure 3C)**, and the relative abundance from each run showed ∼25% MAPE (**Figure 3D)**, indicating high accuracy. These results establish the reproducible recovery of eight barcodes in expected ratios across multiple runs.

**Figure 3.**
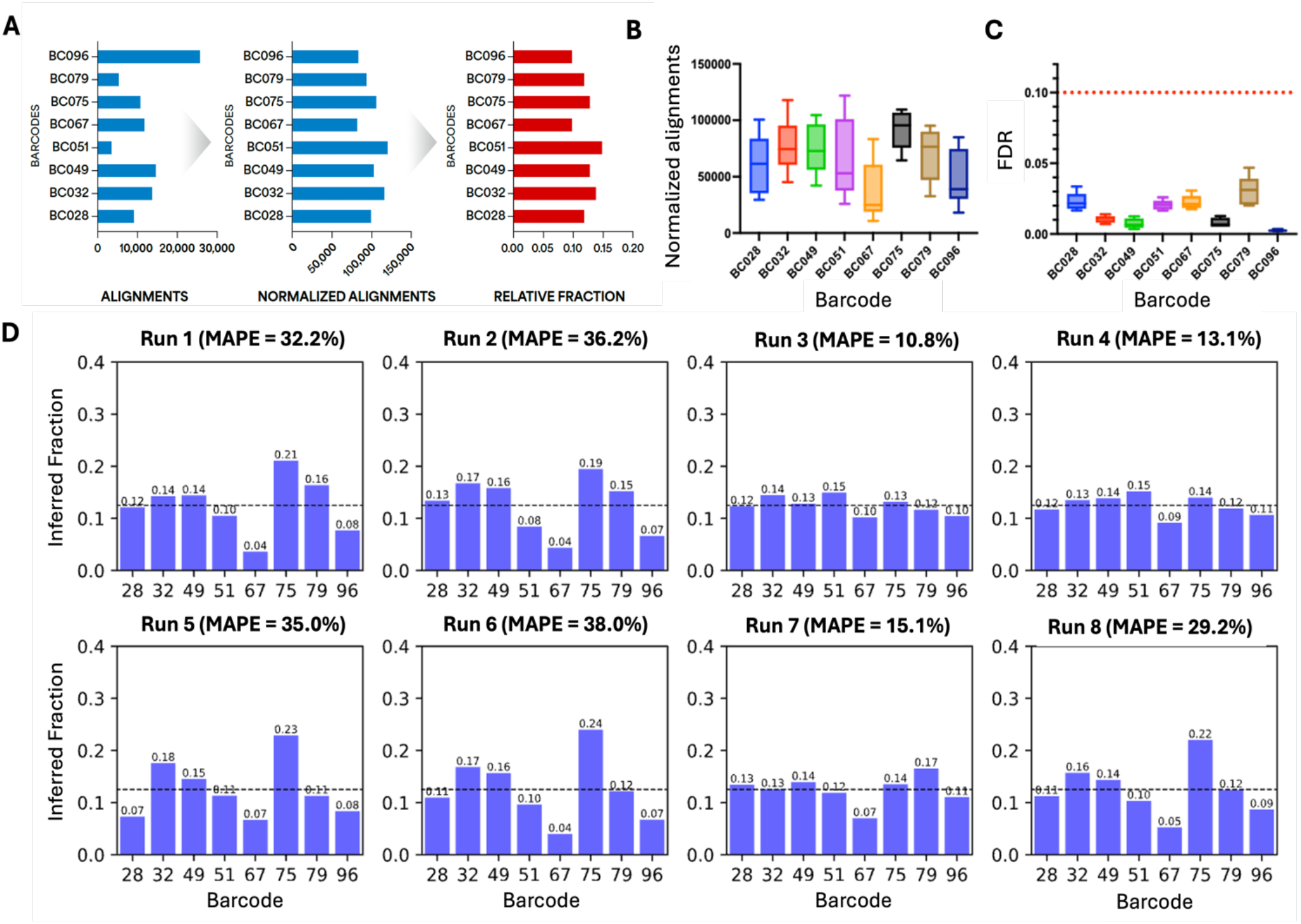
Normalization and reproducibility in 8-barcode mixtures. A) Schematic of normalization workflow showing the strategy for converting raw alignments to normalized alignments, enabling calculation of inferred relative barcode fractions. B) Alignments were normalized and relative fraction recovered for eight runs containing 1:1 eight-barcode mixtures. C) False Discovery Rate (FDR) for normalized alignments across all eight runs; red dotted line indicates 10% FDR. D) Performance summary of recovered inferred fractions for all eight runs plotted individually. MAPE=mean absolute percent error.

### Limit of detection

Next, we tested the limit of detection (LOD) at 25 pmol total input, where each barcode is either at 5 pmol or lower in a plexity of eight. In this experiment, we kept seven barcodes at 1:1 equimolar mix (3.52 pmol each) and varied one barcode by 10-fold lower (0.352 pmol) input concentration. We performed eight total runs, varying one barcode per run to cover all eight barcodes at the lowest input of 352 fmol. We found that four barcodes were successfully recovered (defined as >20 alignments and <10% FDR) at the lowest concentration tested (BC049, BC067, BC075, BC096). However, four barcodes had higher than 10% FDR (BC028, BC032, BC051, BC079) when tested at the lowest input. We then repeated the runs for these four barcodes, increasing the lowest input to 410 fmol. This resulted in successful identification of the four remaining barcodes at <10% FDR (BC028, BC032, BC051, BC079). **Figure 4A** shows an example dataset for the run with BC032 at the lowest input, and **Figure 4B** shows the true fraction plotted against the inferred fraction for this run, resulting in an MAPE of 10.7%. These results demonstrate the relative abundance recovered from the LOD experiment of barcode 32 at the lowest input correlates well with the expected fraction. Likewise, when plotting the expected barcode fraction against the inferred fraction across all eight LOD runs, the calculated cumulative MAPE was 30%. Therefore, the LOD for all eight barcodes were determined to be 410 fmol or below (**Figure 4C)**.

**Figure 4.**
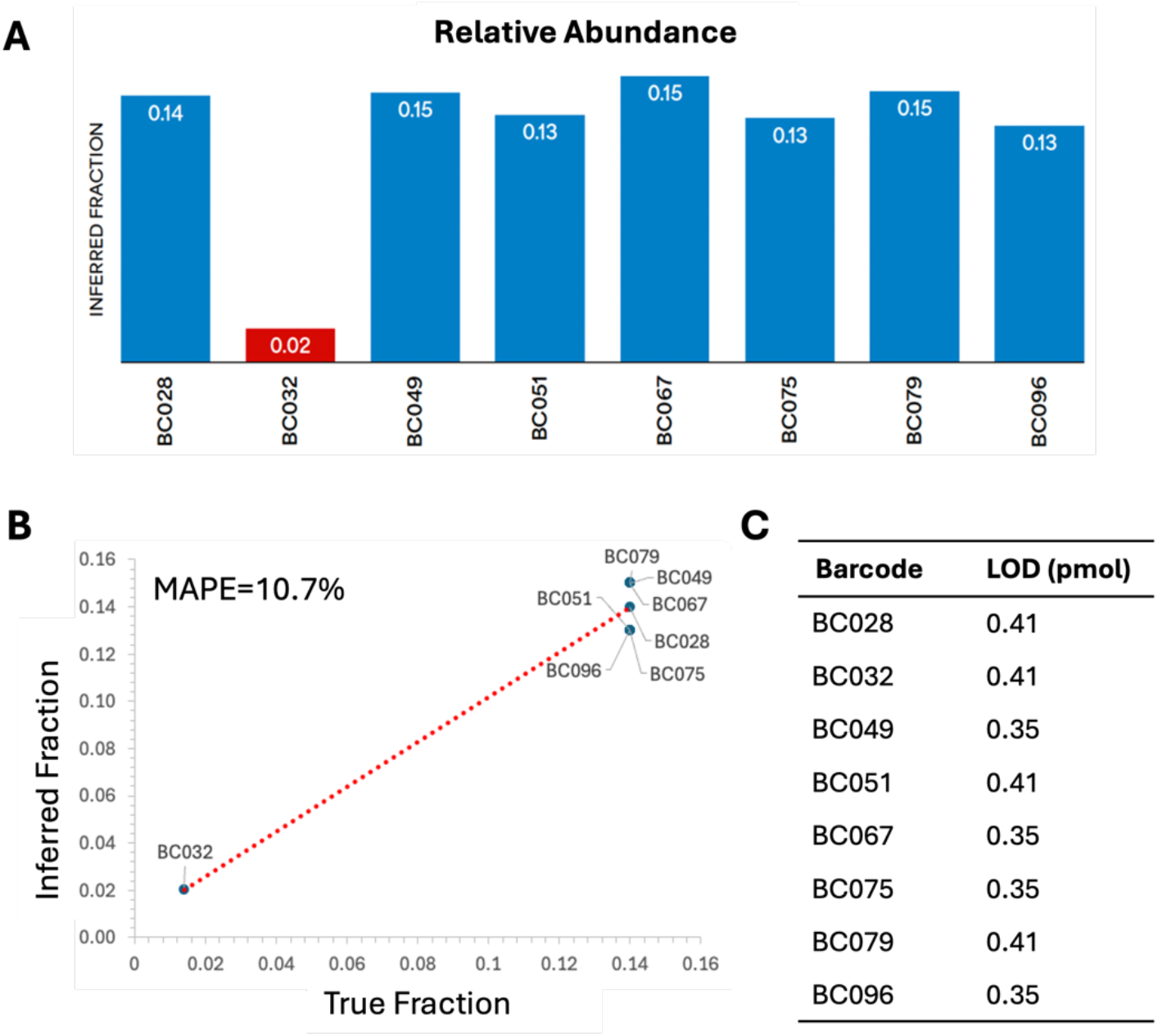
Limit of detection (LOD) for all eight tested barcodes. A) Alignments were normalized for a 10-fold dynamic range titration; in this example, the least abundant barcode (BC032) was positively identified at ∼400 fmol input. B) Inferred fraction vs. true fraction for the data in 4A. C) LOD values in an eight-plex mixture for each barcode tested in this study.

### Dynamic range

We next set out to determine the dynamic range of barcode concentrations measurable within an eight-barcode mixture. We produced 10-fold dynamic ranges by randomly mixing barcodes at 1x (BC051), 0.75x (BC028, and BC096), 0.5x (BC075, and BC079), 0.25x (BC032, and BC049), and 0.1x (BC067). As shown in **Figure 5A**, we identified all barcodes with an FDR < 10%, and the recovered relative abundance shows a good linear correlation after normalization, with an R^2^ of 0.9 and MAPE of 13.9% (**Figure 5B)**. Next, we scrambled these ratios within the same 10-fold dynamic range and performed three different mixes with different barcodes at the 0.1x level (BC032, BC049, and BC075); all mixes with three repeats resulted in successful sequencing as shown in **Figure S6**, with calculated MAPE of 24.3%, 15.9%, and 22.9%, respectively.

**Figure 5.**
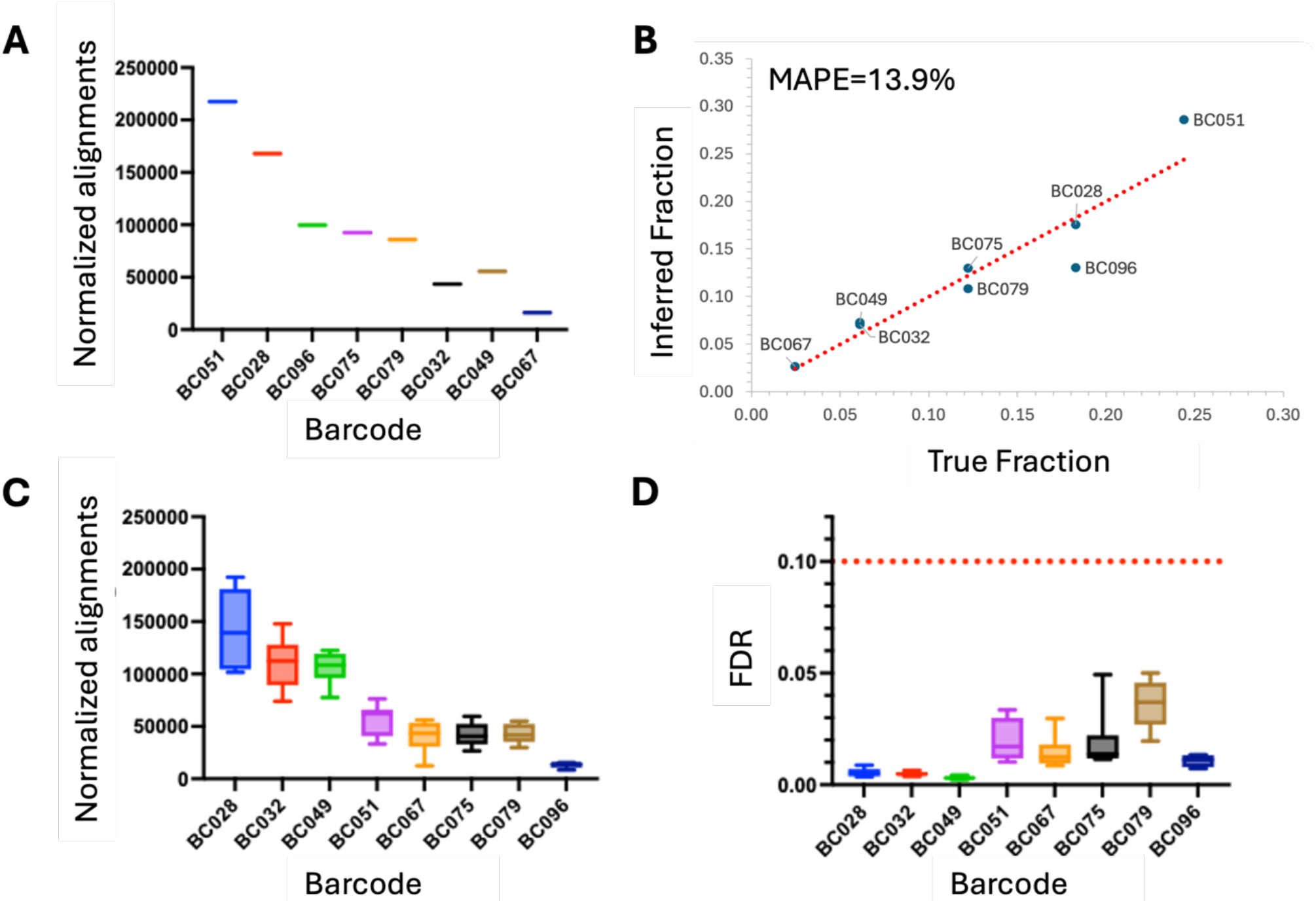
Ten-fold dynamic range of eight barcodes. A) Alignments were normalized for a ten-fold dynamic range titration at the following levels: 1x (BC051), 0.75x (BC028, and BC096), 0.5x (BC075, and BC079), 0.25x (BC032, and BC049), and 0.1x(BC067). B) Inferred fraction vs. true fraction for the data in (A). C) Normalized alignments for eight runs at the following titration levels: 1x (BC028), 0.75x (BC032, and BC049), 0.5x (BC051, and BC067), 0.25x (BC075, and BC079), and 0.1x (BC096). D) FDR for the same runs shown in (C); red dotted line indicates 10% FDR cutoff.

We then tested the reproducibility and robustness of our approach in recovering an unknown dilution within a 10-fold dynamic range. We performed eight additional runs for a 10-fold dynamic range with barcodes at 1x (BC028), 0.75x (BC032, and BC049), 0.5x (BC051, and BC067), 0.25x (BC075, and BC079), and 0.1x (BC096), as shown in **Figure 5C** for the normalized alignments and **Figure 5D** for the plots of FDR for each barcode. These results show that all eight barcodes were successfully identified across all runs with FDRs <10%. In addition, the recovered relative abundance plotted against the true expected fraction from each run showed an MAPE of 21.5%. These results indicate that with an eight-plex barcode mixture with a total input of 25 pmole and the lowest concentration barcode at ∼500 fmol, all eight barcodes are recovered across a 10-fold dynamic range that is still within the LOD. These results demonstrate the robustness of the assay and workflow across a wide range of relative abundances.

### Performance on a mixture of five proteins

Finally, we sought to test the performance of the barcoding workflow in the context of full-length protein expression. We generated five barcoded protein constructs, as shown in **Figure 6A**. These five (IFNg-BC032, PTEN-BC049, TAU441-BC051, UCHL1-BC075, and p53-BC096) were all individually expressed and purified, and the purified barcoded proteins were mixed at 1:1 equimolar ratios (5 pmol per barcoded protein, for a total of 25 pmol) and subjected to the same purification and sequencing workflow as the synthetic barcodes. We prepared eight libraries of this five-protein mix to test the robustness of assay across two lots of Barcoding Kit, two lots of sequencing kits, four lots of chips, four different Platinum instruments, and two operators. The normalized alignments for all eight runs of five-protein mixes show positive identification of all five barcodes (**Figure 6B**), with FDR less than 10% (**Figure 6C**). Across all eight runs, the MAPE ranged from 2.0% to 38.4%, with an average of 16.7% (**Figure 6D**).

**Figure 6.**
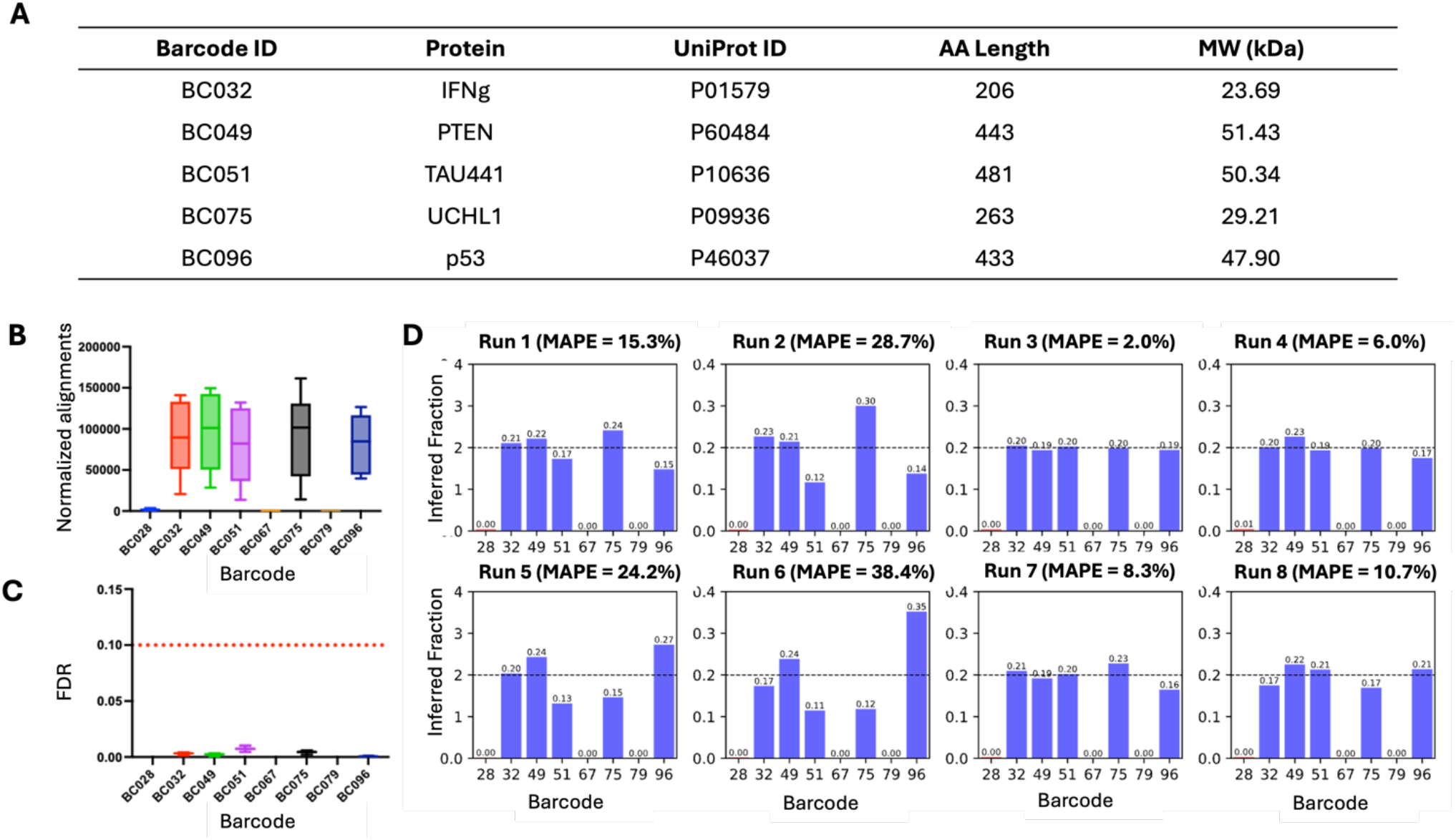
Equimolar mix of five barcoded proteins. A) Summary and characteristics of the five proteins tested in this study. MW = Molecular Weight. B) Normalized alignments recovered across eight runs containing the five proteins mixed at equimolar concentrations. C) FDR across the eight runs shown in (B); red dotted line indicates 10% FDR cutoff. D) Performance summary of recovered inferred fractions for all eight runs plotted individually.

These results demonstrate that the barcoding approach can accurately recover relative abundances in a mixture of full-length proteins.

## Discussion

In this study, we successfully designed and tested a set of barcode constructs for efficient protein labeling and subsequent protein sequencing. Overall, we conducted over 100 protein sequencing runs on over 50 chips, including 10 different lots of sequencing chips, 5 different lots of sequencing reagent kits, and 2 different barcoding kits, all producing an overall MAPE of 24.4% with 95% confidence interval (CI). Over a thousand barcode sequences were generated as part of this effort, with eight optimized peptide sequences chosen for subsequent validation. These barcodes were coupled with affinity tags, flexible linkers, LysC cleavage sites, and sortase tags to enhance barcode enrichment, reduce folding issues, and ensure effective isolation and labeling of proteins. Optimization of the expression construct design also reduced the sample input requirement by 10,000-fold (500 pmol to 50 fmol) and the hands-on time to less than one hour.

Our analysis of barcode normalization and plexity showed successful sequencing and quantification across a 10-fold dynamic range, with relative abundances recovered with high accuracy (MAPE < 25%) across multiple runs. In testing the limit of detection (LOD), barcodes as low as 352 fmol input were identifiable in an eight-plex mixture, and 50 fmol for single proteins. Additionally, when applying this system to a mixture of five proteins expressed in *E. coli*, all proteins were successfully identified with FDR < 10% and a MAPE of 16.7%. These results validate the robustness, accuracy, and sensitivity of the barcoding system for multiplexed proteomics applications.

The ability to accurately normalize barcode abundance across a tenfold dynamic range and detect barcodes at low concentrations (down to 50 fmol) aligns with the need for sensitive, quantitative protein analysis in a variety of applications. However, to achieve successful recovery of relative abundance with high accuracy, it is critical to design experiments that balance the sample input, plexity, and dynamic range, all of which impact the LOD. The sample input directly correlates with the plexity and dynamic range, which then determines the relative fraction of barcodes from lowest to highest abundance. Several key factors can influence sample input, including host expression system, localization, and the target protein. An increase in plexity results in reduced dynamic range, which then requires increased sample input. For example, we hypothesize that a 100-fold dynamic range could be achieved with increased sample input (e.g., 50 pmol or higher). Therefore, it is necessary to consider all of these factors to achieve the target LOD for a given experiment.

Protein barcoding with NGPS has the potential to overcome several limitations of traditional protein analysis methods, such as mass spectrometry and direct labeling. By leveraging the power of NGPS on Platinum for single-molecule resolution, our approach enables precise detection and quantification of protein variants without the need for expensive equipment. In addition, the ability to monitor protein behavior and interactions with minimal disruption to native protein function (due to the compact size of the affinity/barcode tags) is particularly valuable in complex biological systems. Our findings also support the growing role of protein barcoding in applications like nucleic acid therapy development, where direct tracking of protein delivery and function is essential. By ensuring high-fidelity protein sequencing with a broad dynamic range, this work demonstrates how protein barcoding, when paired with NGPS, offers a versatile, scalable, and accessible solution for advancing protein characterization and functional screening.

There are several limitations to this protein barcoding approach that warrant consideration. One key challenge is the potential for barcode interference, particularly in highly complex biological samples where overlapping or similar peptide sequences could lead to cross-reactivity or inaccurate identification. While the use of a carefully optimized set of barcodes with distinct sequences (such as the eight validated in this study) minimizes this risk, the scalability of this method may be impacted when increasing the number of barcodes or when working with particularly complex proteomes. Additionally, while our approach demonstrated high sensitivity and low detection limits in vitro, achieving successful sequencing at extremely low concentrations (i.e., below 50 fmol) still requires careful optimization of sample preparation protocols and sequencing conditions. Lastly, while protein barcoding enables precise tracking of protein identity and abundance, it still requires careful validation in diverse experimental contexts to ensure that barcode incorporation does not affect the native function or interactions of the protein(s) of interest.

Looking ahead, one of the key areas for future development is the scaling up of barcode numbers to enable more complex and diverse proteomic analyses. Expanding the barcode library to include hundreds or even thousands of unique peptide sequences could significantly enhance the versatility of this approach, enabling high-throughput screening and the ability to track a larger number of proteins or protein variants simultaneously. This will require continued development and validation of barcode design to ensure minimal cross-reactivity and false discovery rates as the complexity of the library increases. Additionally, incorporating advanced computational tools for data analysis and barcode normalization will be essential to handle the higher multiplexity and the increased amount of sequencing data. Another promising direction is the demonstration of protein barcoding in vivo. While our current work focuses on in vitro systems, applying this technology in living organisms presents exciting opportunities to track protein behavior, localization, and interactions within physiological contexts. Combining protein barcoding with tissue-specific expression systems could provide insights into protein dynamics in disease models, drug discovery, and gene therapy applications. Ultimately, these advancements will broaden the scope of protein barcoding, making it a powerful tool for both basic research and translational studies in diverse biological and clinical settings.

## Author Contributions

Conceptualization: MC, and JV; Visualization: MC, and JV; Methodology: MC, HH, SH, DP, MR, GHM, MM, IC, and JV; Data Curation: MC, IC, and JV; Investigation: MC, HH, SH, DP, MR, FD, GHM, MM, IC, and JV; Validation: MC, SS, DO, MM, IC, and JV; Formal Analysis: MC, DP, MM, IC, and JV; Writing-Review and Editing: MC, HH, SH, DP, MR, MM, IC, MLC, and JV; Writing-Original: MC, MLC, and JV; Project Administration: MC, and JV; Supervision: JV

## Declaration of Interests

All authors except GHM are employees and shareholders of Quantum-Si, Inc.

## Supplemental Information

Document S1. Figures S1–S6 and Table S1

## Abbreviations

NGPS: Next-Generation Protein Sequencing
LNPs: lipid nanoparticles
ROS: recognizer-ordered sequences
NAA: N-terminal amino acid
MAPE: mean absolute percent error
LOD: limit of detection
FDR: false discovery rate

## Notes

### Summary of Updates

Added author (GHM) who was inadvertently omitted upon first submission.

## References

1. Miyamoto, K., Aoki, W., Ohtani, Y., Miura, N., Aburaya, S., Matsuzaki, Y., Kajiwara, K., Kitagawa, Y., and Ueda, M. (2019). Peptide barcoding for establishment of new types of genotype-phenotype linkages. PLoS One 14, e0215993. 10.1371/journal.pone.0215993.

2. Rossler, S.L., Grob, N.M., Buchwald, S.L., and Pentelute, B.L. (2023). Abiotic peptides as carriers of information for the encoding of small-molecule library synthesis. Science 379, 939–945. 10.1126/science.adf1354.

3. Egloff, P., Zimmermann, I., Arnold, F.M., Hutter, C.A.J., Morger, D., Opitz, L., Poveda, L., Keserue, H.A., Panse, C., Roschitzki, B., and Seeger, M.A. (2019). Engineered peptide barcodes for in-depth analyses of binding protein libraries. Nat Methods 16, 421–428. 10.1038/s41592-019-0389-8.

4. Kumano, S., Tanaka, K., Akahori, R., Yanagiya, A., and Nojima, A. (2024). Using peptide barcodes for simultaneous profiling of protein expression from mRNA. Rapid Commun Mass Spectrom 38, e9867. 10.1002/rcm.9867.

5. Rhym, L.H., Manan, R.S., Koller, A., Stephanie, G., and Anderson, D.G. (2023). Peptide-encoding mRNA barcodes for the high-throughput in vivo screening of libraries of lipid nanoparticles for mRNA delivery. Nat Biomed Eng 7, 901–910. 10.1038/s41551-023-01030-4.

6. Miyazaki, T., Aoki, W., Koike, N., Sato, T., and Ueda, M. (2023). Application of peptide barcoding to obtain high-affinity anti-PD-1 nanobodies. J Biosci Bioeng 136, 173–181. 10.1016/j.jbiosc.2023.07.002.

7. Odunze, U., Rustogi, N., Devine, P., Miller, L., Pereira, S., Vashist, S., Snijder, H.J., Corkill, D., Sabirsh, A., Douthwaite, J., et al. (2024). RNA encoded peptide barcodes enable efficient in vivo screening of RNA delivery systems. Nucleic Acids Res 52, 9384–9396. 10.1093/nar/gkae648.

8. Matsuzaki, Y., Aoki, W., Miyazaki, T., Aburaya, S., Ohtani, Y., Kajiwara, K., Koike, N., Minakuchi, H., Miura, N., Kadonosono, T., and Ueda, M. (2021). Peptide barcoding for one-pot evaluation of sequence-function relationships of nanobodies. Sci Rep 11, 21516. 10.1038/s41598-021-01019-6.

9. Pahl, V., Lubrano, P., Trossmann, F., Petras, D., and Link, H. (2024). Intact protein barcoding enables one-shot identification of CRISPRi strains and their metabolic state. Cell Rep Methods 4, 100908. 10.1016/j.crmeth.2024.100908.

10. Gu, L., Li, C., Aach, J., Hill, D.E., Vidal, M., and Church, G.M. (2014). Multiplex single-molecule interaction profiling of DNA-barcoded proteins. Nature 515, 554–557. 10.1038/nature13761.

11. Vinogradov, A.A., Gates, Z.P., Zhang, C., Quartararo, A.J., Halloran, K.H., and Pentelute, B.L. (2017). Library Design-Facilitated High-Throughput Sequencing of Synthetic Peptide Libraries. ACS Comb Sci 19, 694–701. 10.1021/acscombsci.7b00109.

12. Reed, B.D., Meyer, M.J., Abramzon, V., Ad, O., Ad, O., Adcock, P., Ahmad, F.R., Alppay, G., Ball, J.A., Beach, J., et al. (2022). Real-time dynamic single-molecule protein sequencing on an integrated semiconductor device. Science 378, 186–192. 10.1126/science.abo7651.

13. Sittipongpittaya, N., Skinner, K.A., Jeffery, E.D., Watts, E.F., and Sheynkman, G.M. (2024). Protein sequencing with single amino acid resolution discerns peptides that discriminate tropomyosin proteoforms. bioRxiv, 2024.2011.2004.621980. 10.1101/2024.11.04.621980.

14. Guimaraes, P.P.G., Zhang, R., Spektor, R., Tan, M., Chung, A., Billingsley, M.M., El-Mayta, R., Riley, R.S., Wang, L., Wilson, J.M., and Mitchell, M.J. (2019). Ionizable lipid nanoparticles encapsulating barcoded mRNA for accelerated in vivo delivery screening. J Control Release 316, 404–417. 10.1016/j.jconrel.2019.10.028.

15. Zhang, X., Huang, Y., Yang, Y., Wang, Q.E., and Li, L. (2024). Advancements in prospective single-cell lineage barcoding and their applications in research. Genome Res 34, 2147–2162. 10.1101/gr.278944.124.

16. Gaisser, K.D., Skloss, S.N., Brettner, L.M., Paleologu, L., Roco, C.M., Rosenberg, A.B., Hirano, M., DePaolo, R.W., Seelig, G., and Kuchina, A. (2024). High-throughput single-cell transcriptomics of bacteria using combinatorial barcoding. Nat Protoc 19, 3048–3084. 10.1038/s41596-024-01007-w.

17. Baryshev, A., La Fleur, A., Groves, B., Michel, C., Baker, D., Ljubetic, A., and Seelig, G. (2024). Massively parallel measurement of protein-protein interactions by sequencing using MP3-seq. Nat Chem Biol 20, 1514–1523. 10.1038/s41589-024-01718-x.

18. Mutalik, V.K., Novichkov, P.S., Price, M.N., Owens, T.K., Callaghan, M., Carim, S., Deutschbauer, A.M., and Arkin, A.P. (2019). Dual-barcoded shotgun expression library sequencing for high-throughput characterization of functional traits in bacteria. Nat Commun 10, 308. 10.1038/s41467-018-08177-8.

19. Biggs, B.W., Price, M.N., Lai, D., Escobedo, J., Fortanel, Y., Huang, Y.Y., Kim, K., Trotter, V.V., Kuehl, J.V., Lui, L.M., et al. (2024). High-throughput protein characterization by complementation using DNA barcoded fragment libraries. Mol Syst Biol 20, 1207–1229. 10.1038/s44320-024-00068-z.

20. Chen, I., Dorr, B.M., and Liu, D.R. (2011). A general strategy for the evolution of bond-forming enzymes using yeast display. Proc Natl Acad Sci U S A 108, 11399–11404. 10.1073/pnas.1101046108.

