## Supplementary material for "Protein Barcoding and Next-Generation Protein Sequencing for Multiplexed Protein Selection, Analysis, and Tracking": Document S1

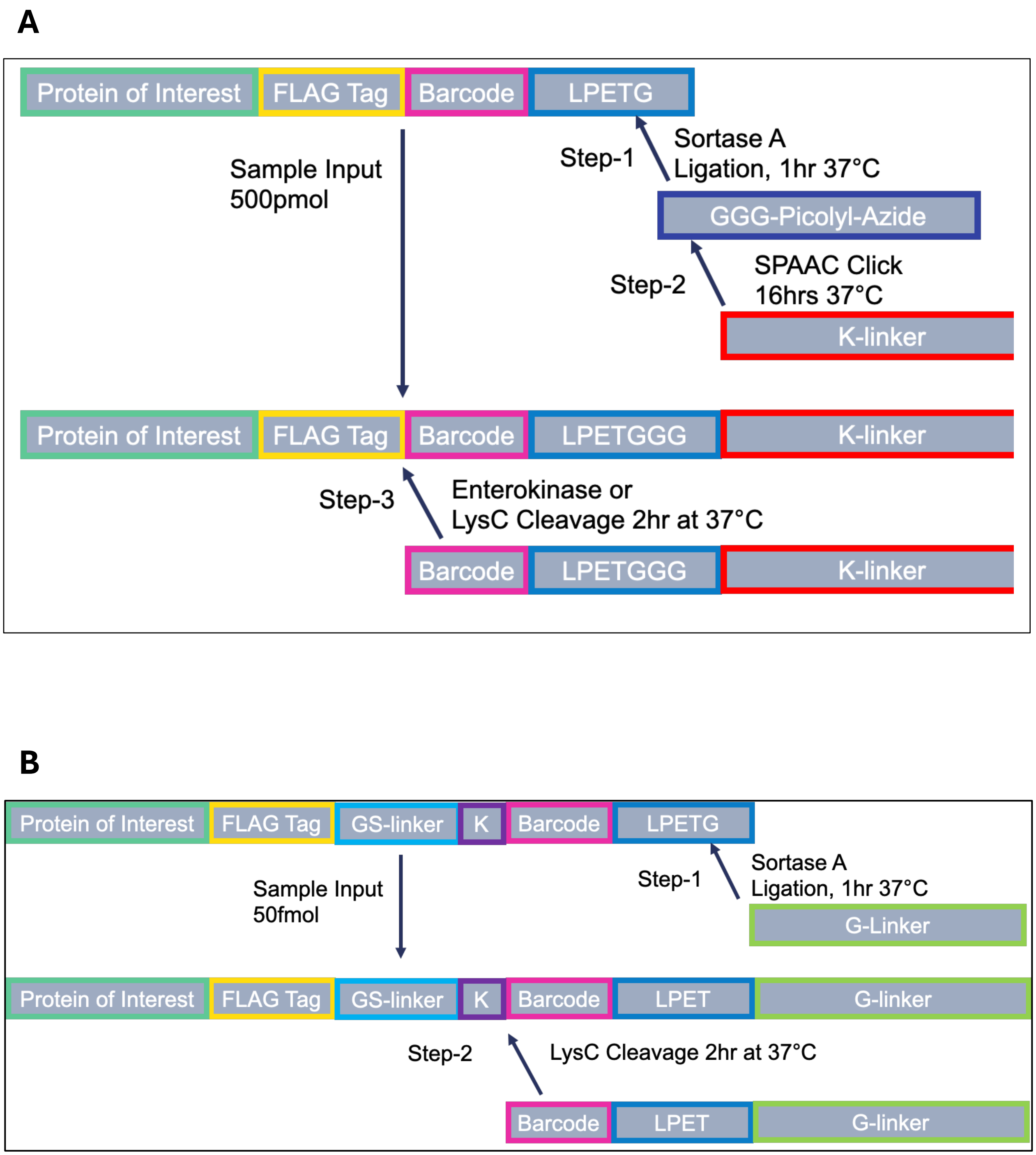


**Figure S1: Overview of Workflow A (A) and Workflow B (B) for barcode expression and purification.** Workflow B, which is shorter and requires a lower sample input, was the final workflow chosen for subsequent experiments.

**A**

| Barcode ID | Protein | UniProt ID | AA | MW, kDa | Sequence |
|---|---|---|---|---|---|
| BC032 | IFNg | P01579 | 206 | 23.69 | MKYTSYILAFQLCIVLGSLGCYCQDPYVKEAENLKKYFNAGHSDVADNGTLFLGILKNWKEESDRKIMQSQIVSFYF<br>KLFKNFKDDQSIQKSVETIKEDMNVKFFNSNKKKRDDFEKLTNYSVTDLNVQRKAIHELIQVMAELSPAAKTGKRKRS<br>QMLFRGRRASQDYKDDDDKGGGGSGGGGSKRQAELFRDYSLPETGHHHHHH |
| BC049 | PTEN | P60484 | 443 | 51.43 | MTAIIKEIVSRNKRRYQEDGFDLDLTYIYPNIIAMGFPAERLEGVYRNNIDDVVRFLDSKHKNHYKIYNLCAERHYDTA<br>KFNCRVAQYPFEDHNPPQLELIKPFCEDLDQWLSEDDNHVAAIHCKAGKGRTGVMICAYLLHRGKFLKAQEALDF<br>YGEVRTRDKKGVTIPSQRRYVYYYSYLLKNHLDYRPVALLFHKMMFETIPMFSGGTCNPQFVVCQLKVKIYSSNSGP<br>TRREDKFMYFEFPQPLPVCGDIKVEFFHKQNKMLKKDKMFHFWVNTFFIPGPEETSEKVENGSLCDQEIDSICSIER<br>ADNDKEYLVLTLTKNDLDKANKDKANRYFSPNFKVKLYFTKTVEEPSNPEASSSTSVTPDVSDNEPDHYRYSDTTDS<br>DPENEPFDEDQHTQITKVDYKDDDDKGGGGSGGGGSKFQRLAELEQALPETGHHHHHH |
| BC051 | TAU441 | P10636 | 481 | 50.34 | MAEPRQEFEVMEDHAGTYGLGDRKDQGGYTMHQDQEGDTDAGLKESPLQTPTEDGSEEPGSETSDAKSTPTAED<br>VTAPLVDEGAPGKQAAAQPHTEIPEGTTAEEAGIGDTPSLEDEAAGHVTQEPESGKVVQEGFLREPGPPGLSHQLM<br>SGMPGAPLLPEGPREATRQPSGTGPEDTEGGRHAPELLKHQLLGDLHQEGPPLKGAGGKERPGSKEEVDEDRDV<br>DESSPQDSPPSKASPAQDGRPPQTAAREATSIPGFPAEGAIPLPVDFLSKVSTEIPASEPDGPSVGRAKGQDAPLEFT<br>FHVEITPNVQKEQAHSEEHLGRAAFPGAPGEGPEARGPSLGEDTKEADLPEPSEKQPAAAPRGKPVSRVPQLKAR<br>MVSKSKDGTGSDDKKAKTSTRSSAKTLKNRPCLSPKHPTPGSSDPLIQPSSPAVCPEPPSSPKYDYKDDDDKGGGG<br>SGGGGSKFALRQDYVAQLPETGHHHHHH |
| BC075 | UCHL1 | P09936 | 263 | 29.21 | MQLKPMEINPEMLNKVLSRLGVAGQWRFVDVLGLEEESLGSVPAPACALLLLFPLTAQHENFRKKQIEELKGQEVS<br>PKVYFMKQTIGNSCGTIGLIHAVANNQDKLGFEDGSVLKQFLSETEKMSPEDRAKCFEKNEAIQAAHDAVAQEGQ<br>CRVDDKVNFHFILFNNVDGHLYELDGRMPFPVNHGASSEDTLLKDAAKVCREFTEREQGEVRFSAVALCKAADYK<br>DDDDKGGGGSGGGGSKNDYRLSQRYLLPETGHHHHHH |
| BC096 | p53 | P46037 | 433 | 47.90 | MEEPQSDPSVEPPLSQETFSDLWKLLPENNVLSPLPSQAMDDLMLSPDDIEQWFTEDPGPDEAPRMPEAAPPVAP<br>APAAPTPAAPAPAPSWPLSSSVPSQKTYQGSYGFRLGFLHSGTAKSVTCTYSPALNKMFCQLAKTCPVQLWVDSTP<br>PPGTRVRAMAIYKQSQHMTEVVRRCPHHERCSDSDGLAPPQHLIRVEGNLRVEYLDDRNTFRHSVVVPYEPPEVG<br>SDCTTIHYNYMCNSSCMGGMNRRPILTIITLEDSSGNLLGRNSFEVRVCACPGRDRRTEEENLRKKGEPHHELPPG<br>STKRALPNNTSSSPQPKKKPLDGEYFTLQIRGRERFEMFRELNEALELKDAQAGKEPGGSRAHSSHLKSKKGQSTSR<br>HKKLMFKTEGPDSDDYKDDDDKGGGGSGGGGSKELFNRALNAFLPETGHHHHHH |

**B**

| Barcode ID | Protein | UniProt ID | AA | MW, kDa | Sequence |
|---|---|---|---|---|---|
| BC-ALQF | SARS-CoV2-RBD | P0DTC2 | 255 | 28.93 | RVQPTESIVRFPNITNLCPFGEVFNATRFASVYAWNRKRISNCVADYSVLYNSASFSTFKCYGVSPTKLNDLCFTNVY<br>ADSFVIRGDEVRQIAPGQTGKIADYNYKLPDDFTGCVIAWNSNNLDSKVGGNYNYLYRLFRKSNLKPFERDISTEIY<br>QAGSTPCNGVEGFNCYFPLQSYGFQPTNGVGYQPYRVVVLSFELLHAPATVCGPKKSTNLVKNKCVNFDYKDDD<br>DKALQFRLFHTDDDLPETGGHHHHHH |
| BC-LFQA | p53 | P46037 | 424 | 47.43 | MEEPQSDPSVEPPLSQETFSDLWKLLPENNVLSPLPSQAMDDLMLSPDDIEQWFTEDPGPDEAPRMPEAAPPVAP<br>APAAPTPAAPAPAPSWPLSSSVPSQKTYQGSYGFRLGFLHSGTAKSVTCTYSPALNKMFCQLAKTCPVQLWVDSTP<br>PPGTRVRAMAIYKQSQHMTEVVRRCPHHERCSDSDGLAPPQHLIRVEGNLRVEYLDDRNTFRHSVVVPYEPPEVG<br>SDCTTIHYNYMCNSSCMGGMNRRPILTIITLEDSSGNLLGRNSFEVRVCACPGRDRRTEEENLRKKGEPHHELPPG<br>STKRALPNNTSSSPQPKKKPLDGEYFTLQIRGRERFEMFRELNEALELKDAQAGKEPGGSRAHSSHLKSKKGQSTSR<br>HKKLMFKTEGPDSDDYKDDDDKLFQARLFHTDDDLPETGHHHHHH |

**Table S1: Summary of protein constructs used in the initial study (development of Workflow A).** MW = molecular weight.

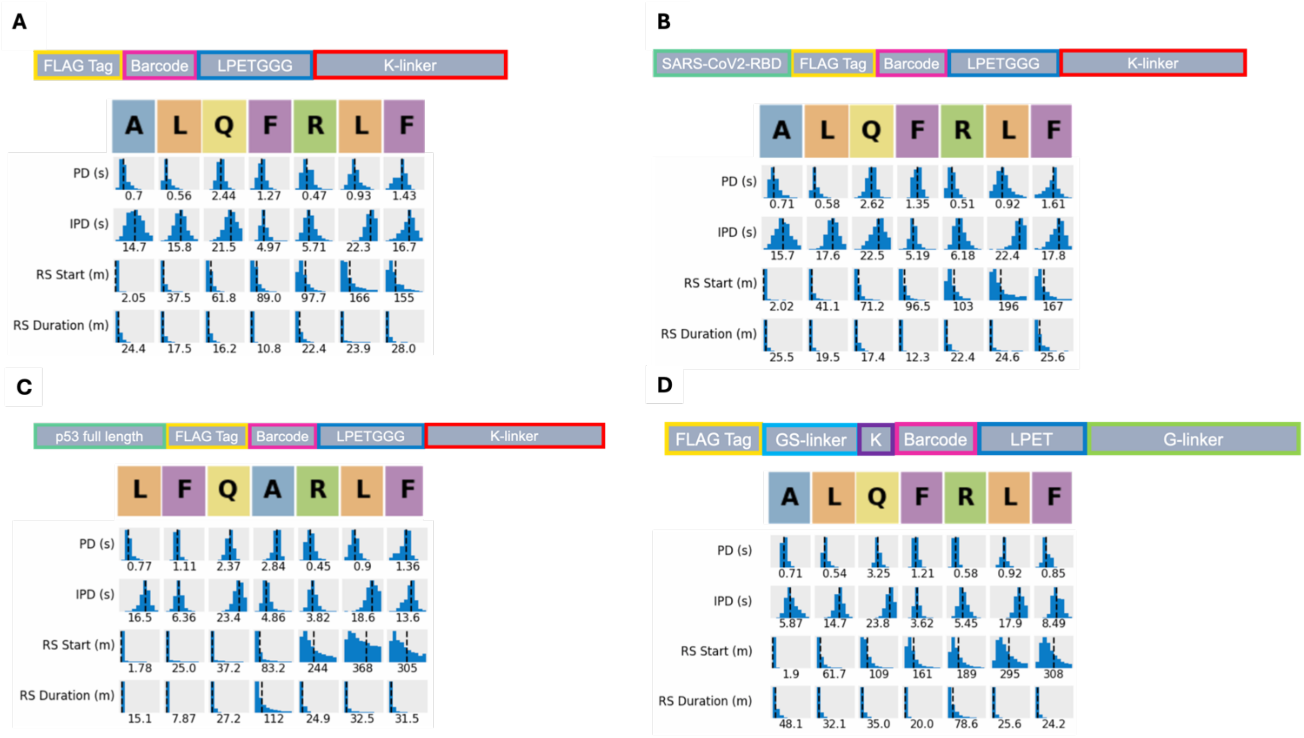


**Figure S2: Kinetics summary for Workflow A with K-linker [(A) BC228, (B) SARS-CoV2-RBD, and (C) p53], and kinetics summary for Workflow B with G-linker [(D) BC265].** PD = pulse duration, IPD = interpulse duration, RS = recognition segment. These kinetic properties are used by the Platinum analysis software to determine the identity and order of amino acids detected during sequencing.

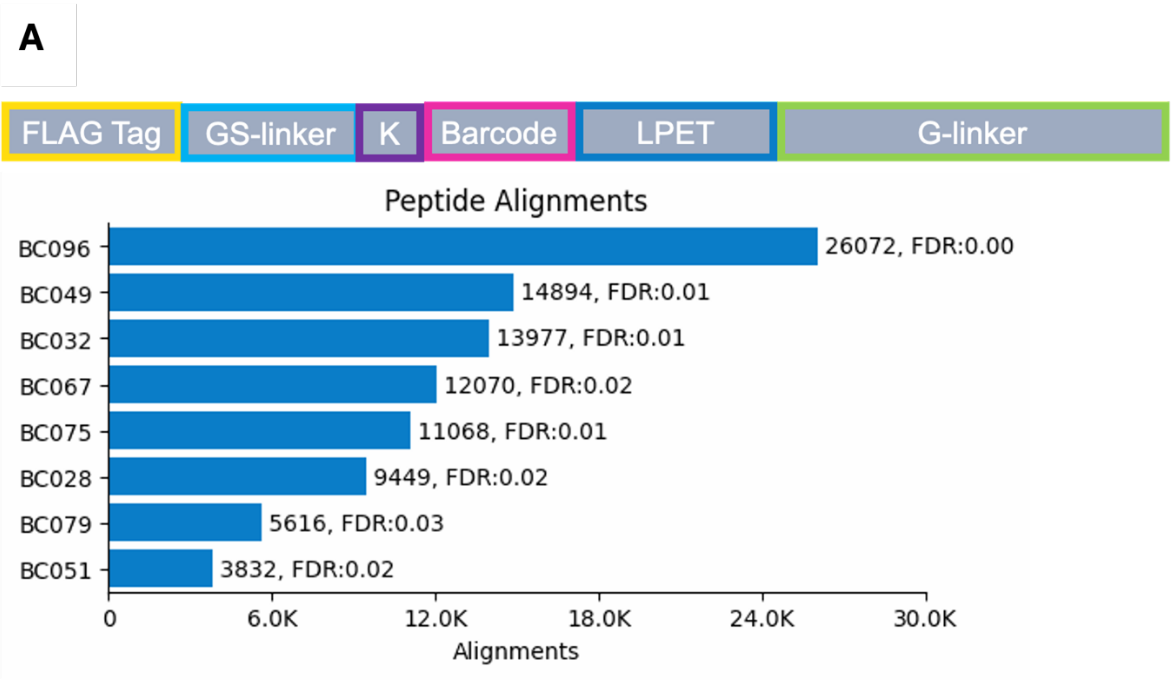


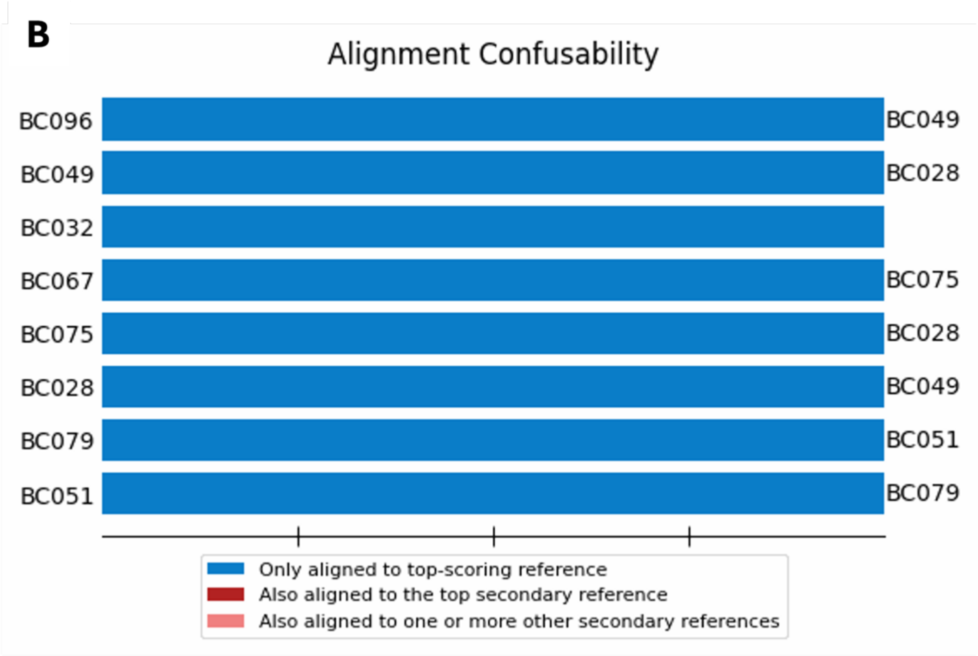


**Figure S3: Alignments (A) and confusability (B) for 1:1 mix of 8 barcodes selected for subsequent validation.** Workflow B construct design with G-linker ligation is also shown in (A). Alignments include read counts and False Discovery Rate (FDR).

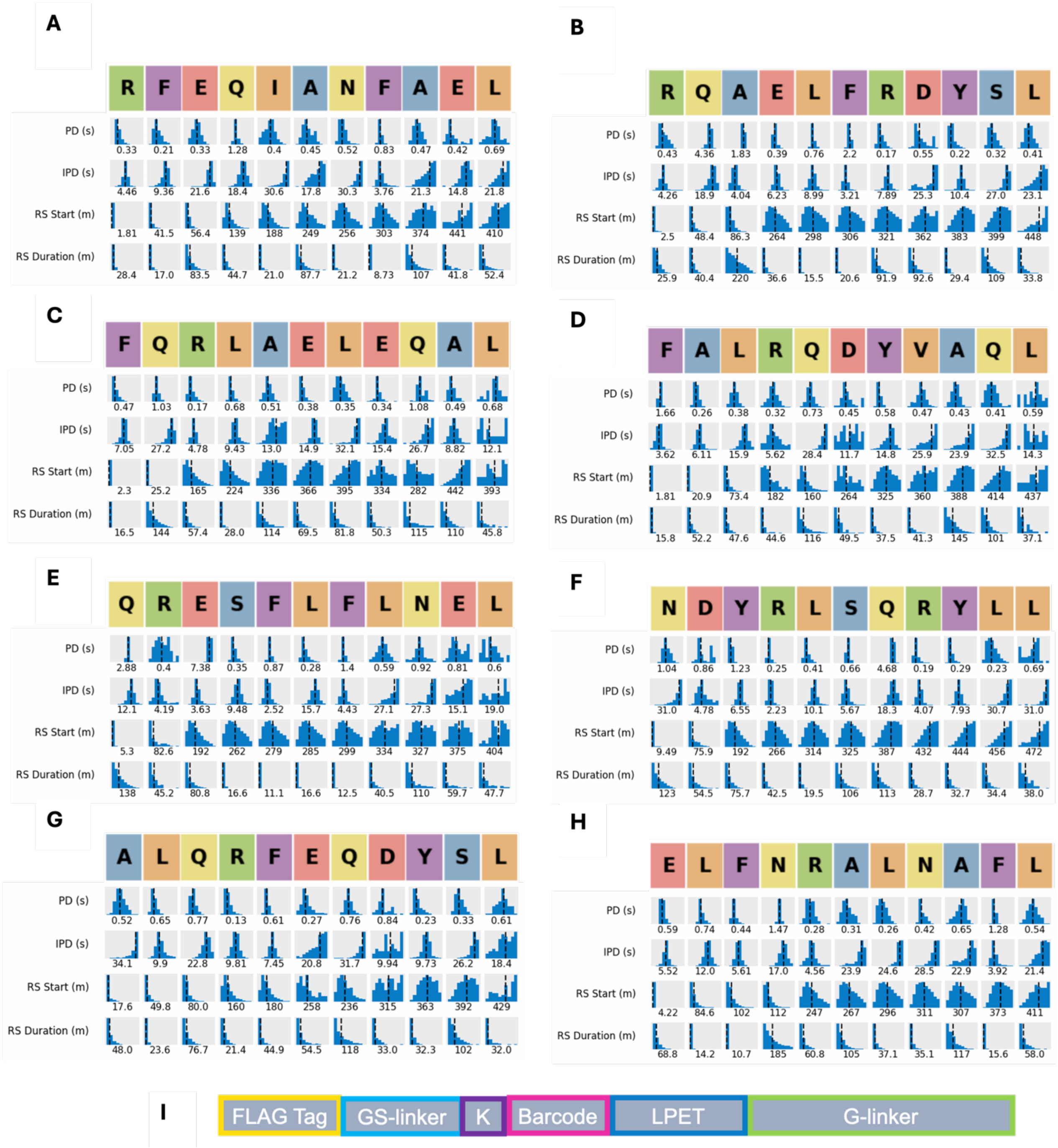

**Figure S4: Kinetics summary plots for 1:1 mix of 8 barcodes. (A) BC028, (B) BC032, (C) BC049, (D) BC051, (E) BC067, (F) BC075, (G) BC079, (H) BC096. (I)** Shows the construct design for Workflow B with G-linker ligation. PD = pulse duration, IPD = interpulse duration, RS = recognition segment. These kinetic properties are used by the Platinum analysis software to determine the identity and order of amino acids detected during sequencing.

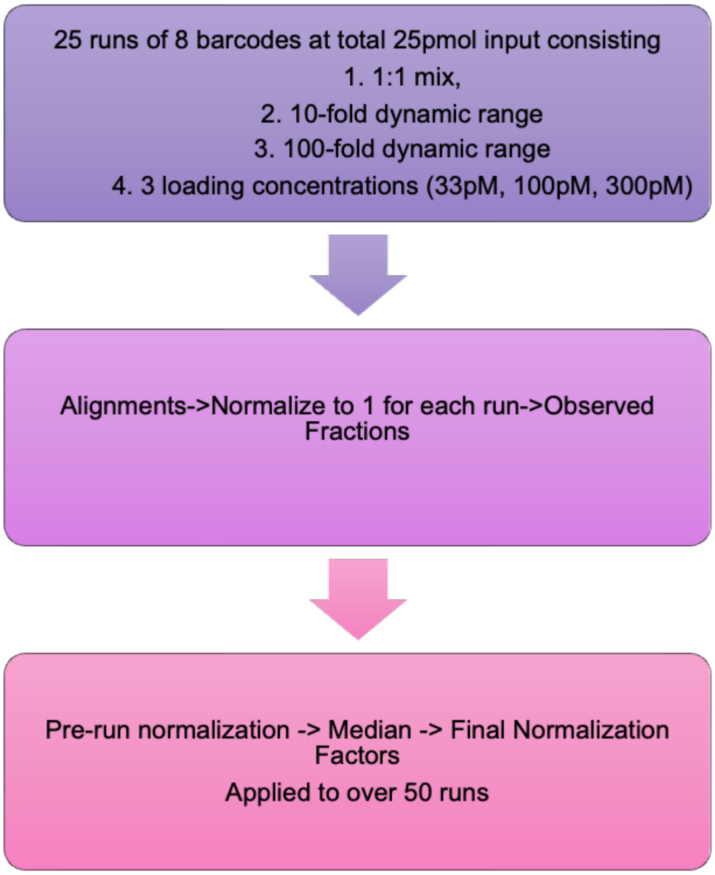


**Figure S5: Overview of workflow to generate barcode normalization factors.** Data from 25 runs is used to generate final normalization factors, which are then applied to subsequent runs to generate inferred fractions for each barcode.

A

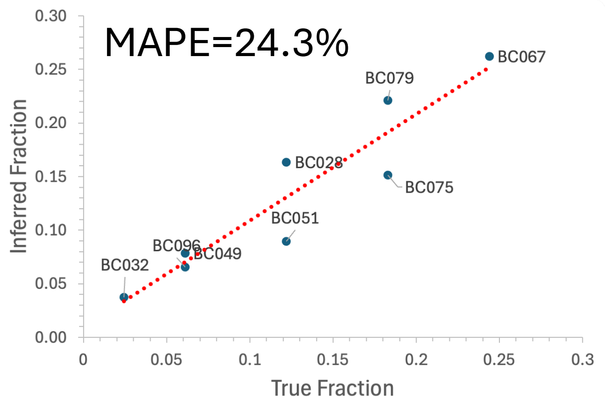


B

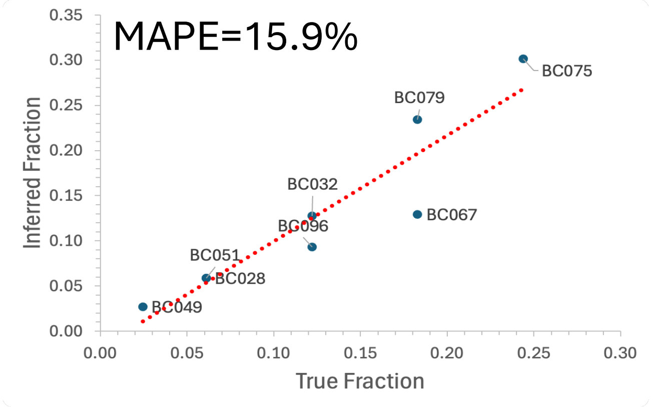


C

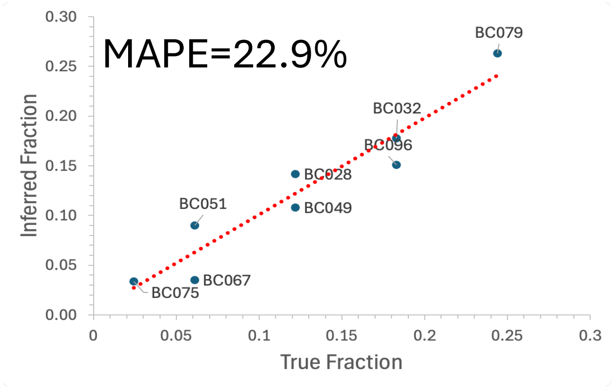


**Figure S6: 10-fold dynamic range data for three different experiments.** Difference between true fraction vs. inferred fraction is shown. (A) BC032 at 10-fold dilution below other 7 barcodes. (B) BC049 at 10-fold dilution below other 7 barcodes. (C) BC075 at 10-fold dilution below other 7 barcodes. MAPE = Mean Absolute Percent Error.
